# Optical Aberration Correction via Phase Diversity and Deep Learning

**DOI:** 10.1101/2020.04.05.026567

**Authors:** Anitha Priya Krishnan, Chinmay Belthangady, Clara Nyby, Merlin Lange, Bin Yang, Loic A. Royer

## Abstract

In modern microscopy imaging systems, optical components are carefully designed to obtain diffraction-limited resolution. However, live imaging of large biological samples rarely attains this limit because of sample induced refractive index inhomogeneities that create unknown temporally variant optical aberrations. Importantly, these aberrations are also spatially variant, thus making it challenging to correct over wide fields of view. Here, we present a framework for deep-learning based wide-field optical aberration sensing and correction. Our model consists of two modules which take in a set of three phase-diverse images and (i) estimate the wavefront phase in terms of its constituent Zernike polynomial coefficients and (ii) perform blind-deconvolution to yield an aberration-free image. First, we demonstrate our framework on simulations that incorporate optical aberrations, spatial variance, and realistic modelling of sensor noise. We find that our blind deconvolution achieves a 2-fold improvement in frequency support compared to input images, and our phase-estimation achieves a coefficient of determination (*r*^2^) of at least 80% when estimating astigmatism, spherical aberration and coma. Second, we show that our results mostly hold for strongly varying spatially-variant aberrations with a 30% resolution improvement. Third, we demonstrate practical usability for light-sheet microscopy: we show a 46% increase in frequency support even in imaging regions affected by detection and illumination scattering.

## 1. Introduction

High-resolution optical imaging is of great importance in fields like astronomy, microscopy, and biomedical imaging. However, the resolution of the acquired images is often limited by system as well as sample-induced optical aberrations. While system aberrations are caused by limitations in optical design, sample-induced aberrations may be caused by complex refractive index changes within the sample that vary in both space and time. One such example is the live imaging of fluorescently labelled embryos using light-sheet microscopy [1, 2]. Given a known and invariant Point Spread Function (PSF), a popular solution for addressing system aberrations is to apply deconvolution algorithms such as Richardson-Lucy (RL) [3] which can be extended for multi-view imaging [4, 5]. If the PSF is unknown, blind deconvolution (BD) algorithms can jointly estimate the image and PSF [6]. Another solution of choice is Adaptive Optics (AO) which implements wavefront sensing (i.e. measures deviation from a planar wavefront) and correction in hardware by means of a Shack-Hartmann sensor and deformable mirror, respectively. While highly effective, corrections afforded by AO are generally only valid over a small field-of-view, or require prohibitively complex hardware setups capable of multi-conjugate correction [7]. In contrast, other phase retrieval approaches can estimate lower order aberrations across a wide field-of-view from multiple images acquired with known phase offsets [8], and can jointly estimate object and aberrations [9].

With the success of Convolutional Neural Networks (CNNs) in performing various image-based tasks (see [10] for a comprehensive review), CNN models have been developed for deconvolution and wavefront sensing. Some of these approaches use a blur kernel that includes focus and astigmatism only or use larger kernels that are not physically realisable in a microscope [11, 12] and typically do not take advantage of additional information from phase-diverse images. Other approaches model the image estimation part with a CNN and use a deconvolution module [11, 13] to extract the PSF, or use existing CNN architectures such as ResNet and Inception to regress Zernike coefficients [12, 14, 15], and use iterative Richardson-Lucy to estimate the image but not both.

Here we present a CNN approach for blind deconvolution and phase estimation, both of which take advantage of phase-diverse image acquisitions. The key aspects of our work are: (i) the extraction of 3D convolutional features from phase-diversity stacks – so as to capture axial extent of PSF and thus facilitate deconvolution; (ii) the estimation of up to 4^*th*^ order Zernike coefficients including astigmatism, coma and trefoil; (iii) the training of an end-to-end framework for joint estimation of the PSF and deconvolved image; (iv) a demonstration of correction on spatially varying aberrations; (v) the application on a phase-diverse dataset acquired on a light-sheet microscope – to better understand the performance and limits of our approach.

### Related Work

Earlier CNN models for deconvolution unroll the iterative algorithm and solve it as a nonlinear regression problem. Zhang *et. al.* developed a cascaded Fully Convolutional Neural Network (FCNN) to address the effect of noise and sensitivity to image-priors in non-blind deconvolution of natural images. The model denoised the vertical and horizontal gradients which were used as image priors to an iterative deconvolution algorithm [11]. Schuler *et.al.* used a similar multistage model for blind deconvolution of natural images with each stage consisting of a feature extraction, kernel estimation, and image estimation module. The resulting model outperformed conventional approaches for small and medium size blur kernels, but failed for larger kernels [13]. Recently, Shajkofci *et. al.* fine-tuned Alex-net and Res-net to estimate optical aberrations in microscopy images, and performed semi-blind deconvolution using total variation *RL* [12]. On a related note, Paine *et.al.* trained an Inception model and its variant for estimating a good initial guess of the wavefront [14]. and Nishizaki *et.al.* used iterative wavefront sensing approaches in combination with an image pre-conditioner for overexposure, defocus, and scatter [15].

## 2. Theory

### Wavefront parameterization

Wavefront phase is commonly modeled as a series expansion of Zernike polynomials [16], which are orthogonal polynomials on the unit circle:

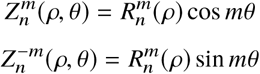

A double indexing scheme is used, where index n represents the highest power of the polynomial for the radial component (ρ: 0 - 1), index m represents the frequency of the azimuthal component(θ: 0 - 2π), and 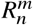 is the radial component of the Zernike polynomials [12]. Wavefront aberrations are described as the sum of Zernike modes *Z*_*j*_ multiplied by their coefficient *C*_*j*_ indexed by the Zernike mode number j:

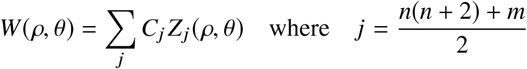

Hence, the Zernike coefficient *vector Z* = [*C*_*j*_] is sufficient to fully characterise the wavefront.

### Image formation

At a focus distance ∊, the point spread function is related to wavefront distortion by:

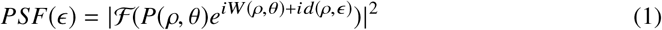

Where, P is the pupil function and ℱ is the Fourier transform. The defocus-only *phase diversity* [8] under paraxial approximation is given by:

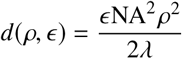

where NA is the numerical aperture, λ is the optical wavelength, and ∊ is the focus distance. For simplicity our model assumes that light is emitted from a single plane within the sample – as if illuminated by an infinitely thin and extended light-sheet. In what follows we consider an objective with a numerical aperture of N A = 0.8 and a pixel size of 0.406 µm.

### Problem statement

The observed *aberrated* image is given by: *I*_0_ = *I* * *PSF* 0 where *I* is the non-aberrated image. In addition to optical aberrations, these images are also degraded by shot noise and electronic noise. For the purpose of phase retrieval, we also acquire two additional slightly out-of-focus *phase-diverse* images: *I*_− 1_ = *I* * *PSF* (− ∊) and *I*_∊1_ = *I* * *PSF* (+∊) where ∊ is the absolute defocus. While more than three phase-diverse images can be used, one should also consider the practicality and cost in time of acquiring more than three images. Our goal is, given the images *I*_0_, *I*_−1_, and *I*_+1_, to obtain an estimate *I* ^′^of the the true object image *I* and the corresponding estimate *Z* ^′^of the true Zernike coefficients *Z*.

### Implicitly learned image prior

Classical iterative solutions [8, 17] for estimating the true object image *I* impose analytical constrains – such as e.g. non-negativity – on the structure of solutions *I* ^′^. The strength of generality of classical – non deep-learning – methods is also a weakness because for specific imaging applications strong solution priors exist that can substantially improve reconstruction quality. One of the key advantages of CNNs is their ability to learn both the inverse function but also an implicit prior on solutions [10, 18]. In this work we focus our attention on light-sheet microscopy images of fluorescently labelled nuclei in developing zebrafish (*D. rerio*) and fruit fly (*D. melanogaster*) embryos. Imaging within such large embryos is challenging because of sample induced aberrations that rapidly degrade image quality for deep imaging planes (>50µm). Yet, these images are highly stereotypical: while exhibiting some variance, cell nuclei have a typical size, shape and spatial distribution, and thus offers an opportunity for improved image deconvolution and phase estimation using deep learning.

## 3. Methods

### Sample preparation

Mounting and imaging of zebrafish embryos were handled in accordance with the University of California San Francisco guidelines and were approved by the Institutional Animal Care and Use Committee. Imaging experiments were performed on the zebrafish transgenic line *h2afva:h2afva-mCherry*. After dechorionation, embryos were mounted in 0.8% low melting point agarose inside a 1.5 mm inner diameter glass capillary. After agarose solidification, the agarose section containing the embryo is extruded from the glass capillary with a plunger. The sample is then placed in a custom-made multi-view light-sheet microscope [2] after filling the imaging chamber with E3 fish medium.

### Microscopy

We use images of developing zebrafish and fruit fly embryos previously acquired on a light-sheet microscope [2] for the simulations described below, as well as newly acquired phase-diverse zebrafish images for evaluation. Imaging is done with Nikon 16 × 0.8*N A* water-dipping objectives and an Orca Flash 4.0 sCMOS camera resulting in a pixel size of 0.406 µm. we illuminate the sample with a 561 nm laser light-sheet and filter detection using a 610 / 75 bandpass filter. Exposure is set to 20 ms.

### Simulated training data

For each dataset (zebrafish and fruit fly), the images are first denoised using Noise2Self [19], then image tiles of size 256 × 256 pixels are randomly selected, and then normalised to have intensity values within [0, 1]. The tiles are convolved with PSF kernels of size 31 × 31 pixels to generate aberrated images for training. For simplicity we assume that fluorescent light is emitted from an infinitely thin slab of material co-planar with the detection plane. To compute the PSF images, we use Eq. 1. To model the sample induced aberrations, Zernike coefficients of orders 2, 3 and 4 are sampled from a uniform random distribution between [−1, 1]. The sampled Zernike coefficients are multiplied by a bi-exponential decay term that adjusts the contributions of the various Zernike orders. We do not include lateral image shifts 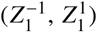 and focus 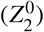 since these can be corrected with registration and auto-focus methods [20]. A defocus term is added and subtracted from these Zernike coefficients to obtain out-of-focus PSFs to generate the phase-diverse images *I*_0_, *I*_− 1_, and *I*_+1_. we use a defocus ∊ of +/ −2µm Poisson noise (*n* = 150) and Gaussian noise (σ = 0.001) are applied to the blurred tiles to simulate both realistic sensor and photon shot noise.

### Model overview

As shown in Fig. 1 our model consists mainly of a deconvolution model *HUnet* and phase estimation model *Znet*. An additional pre-trained differentiable model *PSFnet* is used for generating a PSF image from the predicted Zernike coefficients – this model is used for enforcing consistency between the deconvolution and phase estimation. All models are implemented in python using the Deep-Learning library *PyTorch*. In the following paragraphs we give architectural details on each model.

**Fig. 1.**
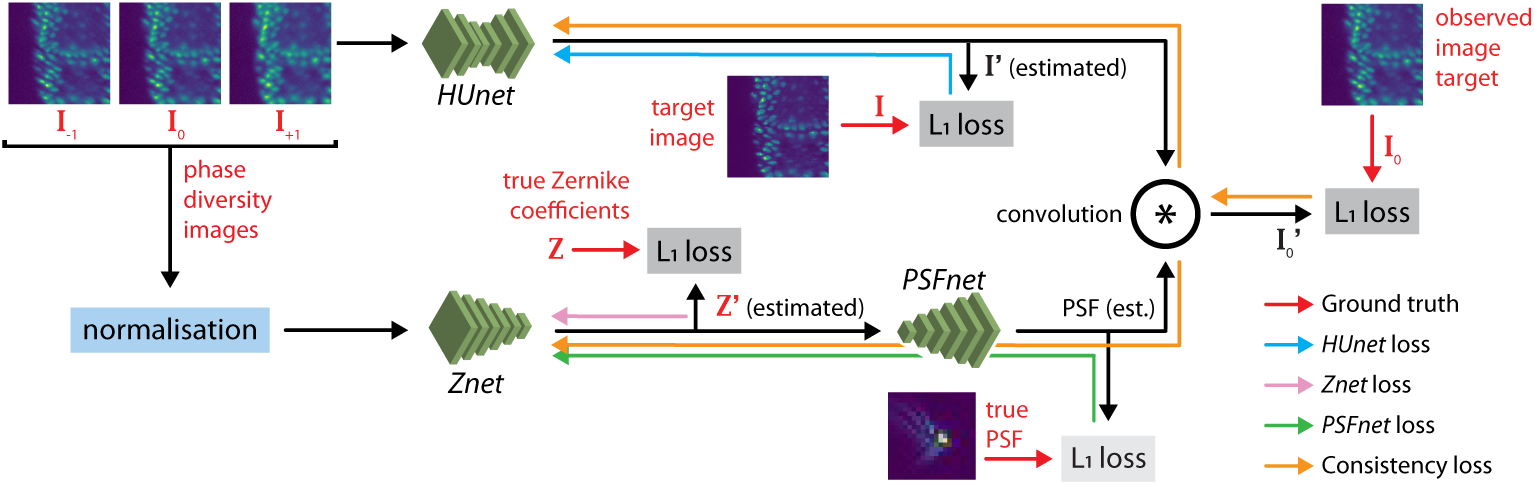
Model Overview. The two networks *HUnet* and *Znet* compute the deconvolved image *I* ^′^ and Zernike coefficients *Z* ^′^, respectively, from phase-diverse images *I*−, *I*_0_, and *I*_+_. *PSFnet* is a differentiable model used for enforcing self-consistency that computes the PSF from a corresponding Zernike vector *Z* ^′^. We show all four losses considered and corresponding gradient paths.

#### Hunet

This network is a 3D-2D hybrid version of the U-net [21], where the down-sampling arm uses 3D convolutions with 3 × 3 × 3 kernels and the up-sampling arm uses 2D convolutions implemented via anisotropic 1 × 3 × 3 kernels (Fig. 2a). The use of 3D convolutions allows the network to identify 3D Patterns within the three images (*I*_−_, *I*_0_, and *I*_+_), which together are equivalent to a defocus stack. Skip connections consist of *N* × 1 × 1 convolutions (with no zero-padding) followed by batch normalisation, where *N* is the number of phase diverse images and batch normalization. We use nearest-neighbour interpolation for up-sampling. This network, with 82 layers and 1.23 million trainable parameters, takes a stack of three images *I*_0_, *I*_−1_, and *I*_+1_ and predicts the deconvolved image *I*’.

**Fig. 2.**
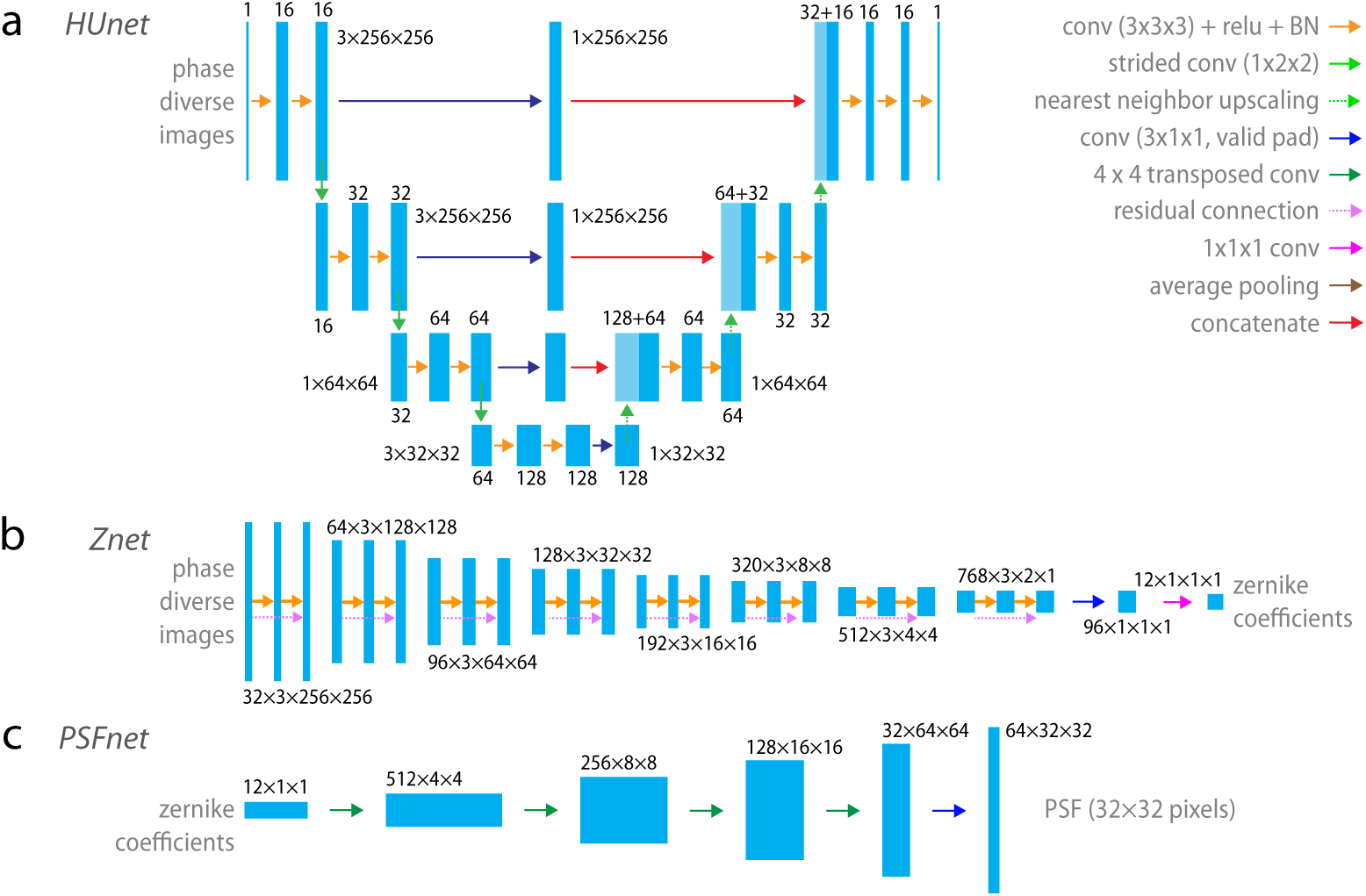
Detailed model architectures for **(a)** *HUnet*, **(b)** *Znet*, and **(c)** *PSFnet. HUnet* is a hybrid 3D-2D UNet [23] that receives three phase diverse images *I*−, *I*_0_, and *I*_+_ as a single stacked 3D image, and returns a single 2D deconvolved image *I* ^′^. *Znet* is made of 8 residual convolutional blocks that take the three phase-diverse images and returns a single vector of 12 Zernike coefficients. *PSFnet* takes a vector of 12 Zernike coefficients and returns the PSF sampled on a 32 × 32 image. Note: these architectures can be easily be adapted to accommodate an arbitrary number *N* of phase diverse inputs.

#### Znet

This network predicts Zernike coefficients *Z* from the three phase-diverse images *I*_0_, *I*_−1_, and *I*_+1_. In a manner similar to the *HUnet*, we use 3D convolutions with 3 × 3 × *N* (*N*: number of phase diverse images, *N* = 3) kernels, down-sampling, and residual connections to convert the phase-diverse input image tensor of size 3×256×256 to an estimated Zernike vector *Z* ^′^ of size 1 × 1 × 12 (see Fig. 2b). The *Znet* has 76 layers and 14.15 million parameters.

#### PSFnet

This network provides a differentiable and numerically stable learned approximation of the function that computes the PSF from estimated Zernike coefficients *Z* ^′^ (see Eq. 1). It consists of four blocks with each block consisting of transposed convolution for up-sampling, batch-normalization, ReLU nonlinearity and dropout layers [22] (Fig. 2c). The final block consists of convolution and hyperbolic tangent activation layers. The network has 13 layers and 2.85 million trainable parameters.

### Joint training

We hypothesise that training both *HUnet* and *Znet* jointly and enforcing consistency between the deconvolved image *I* ^′^ and the estimated Zernike coefficients *Z* ^′^ will encourage the networks to truly learn image deconvolution and Zernike coefficient estimation and possibly prevent the models from over-fitting on image features. Four losses are used for joint training: *L*_1_ loss between *I* and *I* ^′^, between *Z* and *Z* ^′^, between the output of *PSFnet* and the true PSF (*PSFnet* weights are held fixed but gradients can pass through *Znet*), and a consistency loss. The deconvolved image *I* ^′^ from *HUnet* is convolved with the predicted PSF to infer a reconstructed aberrated image 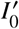 from both *I* ^′^ – computed by *HUnet*, and *Z* ^′^ – computed by *Znet*. The *L*_1_ loss between this predicted blurred image 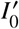 and the center image (*I*_0_) provides the consistency loss that is backpropagated to update parameters of both *HUnet* and *Znet* (Fig. 1).

### Training parameters

*HUnet* and *Znet* are trained using *L*_1_ loss, the Adam optimiser, and a learning rate of 10^−4^. We use a batch size of 16 and the models are trained for 300 epochs with a 70 − 15 − 15 split for train, validation and test sets. To prevent overfitting we use a dropout of 0.2 and use a random subset of tiles for training in every epoch. Training takes 17 hours for *HUnet* and 24 hours for *Znet* (NVIDIA Titan X). *PSFnet* is trained on a separate set of 10000 pairs of Zernike coefficients (sampled in the same manner as previously described for the *HUnet* training data) and analytically computed PSF images of size 31 × 31 pixels. The PSFs are normalized to have an area under the curve of one. The *PSFnet* achieves an SSIM of 0.967 between the target and predicted PSF images.

### Iterative non-blind deconvolution

We compare the performance of our approach against three iterative *non-blind* deconvolution algorithms: (i) Richardson-Lucy Total Variation (RLTV) which augments Richardson-Lucy’s approach [24] with a TV prior [25], (ii) Iterative Constraint Tikhonov-Miller (ICTM) which minimizes a Tikhonov functional and enforces non-negativity at each iteration [26], and (iii) Bounded-Variable Least Squares (BVLS) [27]. For all three approaches we use *DeconvolutionLab2* implementations [27].

### Evaluation metrics

Mean structural similarity (SSIM) [28], mean coefficient of determination (*r*^2^) over the test set are used as metrics for evaluating *HUnet* performance. Mean coefficient of determination and mean absolute error (MAE) for each predicted Zernike coefficient is used for evaluating *Znet* performance.

## 4. Results

### Deconvolution performance

Overall, our blind deconvolution model (*HUnet*) trained on the zebrafish dataset performed slightly better than that trained on fly dataset (SSIM of 0.421 for fly versus 0.932 for zebrafish, see Table 1) which might be attributed to the larger nuclei found in the zebrafish dataset. Importantly, both models generalised well when trained on one dataset and applied onto the other – with models trained on zebrafish generalizing marginally better (Table 2).

**Table 1.** *HUnet* performance and comparison with iterative non-blind deconvolution. PD stands for Phase Diversity.

|  | fly |  | zebrafish |  |
| --- | --- | --- | --- | --- |
| | $r^2$ | SSIM | $r^2$ | SSIM |
| HUnet With PD | **0.929** | **0.421** | **0.987** | **0.932** |
| HUnet No PD | 0.927 | 0.404 | 0.982 | 0.922 |
| ICTM | 0.693 | 0.317 | 0.870 | 0.556 |
| BVLS | 0.662 | 0.294 | 0.856 | 0.512 |
| RLTV | 0.623 | 0.295 | 0.818 | 0.458 |
| in-focus | 0.691 | 0.264 | 0.869 | 0.475 |

**Table 2.** *HUnet* generalisation between fly and zebrafish. Training dataset along rows and testing dataset along columns. Numbers are SSIM.

|  | fly (test) | zebrafish (test) |
| --- | --- | --- |
| fly (train) | 0.421 | 0.727 |
| zebrafish (train) | 0.466 | 0.932 |

### Comparison with state-of-the-art iterative non-blind deconvolution

As stated previously, We compared our phase diversity *blind* deconvolution against classical deconvolution *non-blind* algorithms to which we provide the *true* PSF. Of the three deconvolution methods compared, ICTM performs best with a mean SSIM of 0.317 on the fly dataset and 0.556 on the zebrafish dataset, which is an improvement of approximately 20% and 17% compared to blurry input images for the respective datasets (Table 1). However, our deconvolution model (*HUnet*) achieved a mean SSIM of 0.421 on the fly dataset and of 0.932 on the zebrafish dataset – an improvement of approximately 60% and 95% compared to the blurry input images (Table 2). The better performance of *HUnet* compared to non-blind deconvolution is most likely due to its ability to implicitly learn a strong reconstruction prior from the stereotypy of nuclei images – i.e. from the repetitive features found in nuclei images.

### Frequency domain resolution analysis and comparison

As shown in Fig. 3 our *HUnet* deconvolution model recovers noise-free deconvolved images. Detailed image frequency analysis shows that for low frequencies (red triangle marker, features above 5 pixels) the performance of *HUnet* is comparable to that of the best performing deconvolution (ICTM), but for higher spatial frequencies (features of scale below 5 pixels), *HUnet* outperforms ICTM by rejecting the noise and in-filling lost frequency details with a very natural-looking apodization. In contrast, ICTM restored images show strong artifacts, which as shown in Fig. 3 have their origin in erroneous high frequency restitution (see black triangle marker). Using Mizutani’s single image resolution estimate [29] we can quantify that improvement further: we find that *HUnet* deconvolved images have 2-fold wider frequency support than the best focussed input image: 1.99 and 2.34 for fly and zebrafish, respectively. Despite the artefacts mentioned previously, ICTM deconvolution performs well but still underperforms compared to *HUnet*: 1.52 and 1.89 fold improvement in frequency support for fly and zebrafish, respectively. Paradoxically, and yet as expected, *HUnet* exhibits a wider frequency support than the ground truth itself – this is because of (i) the denoising and frequency in-painting at high frequencies, (ii) the nuclei prior is learned over the whole training set based on real microscopy images with a wide distribution of sharpness which affords opportunities for further sharpening.

**Fig. 3.**
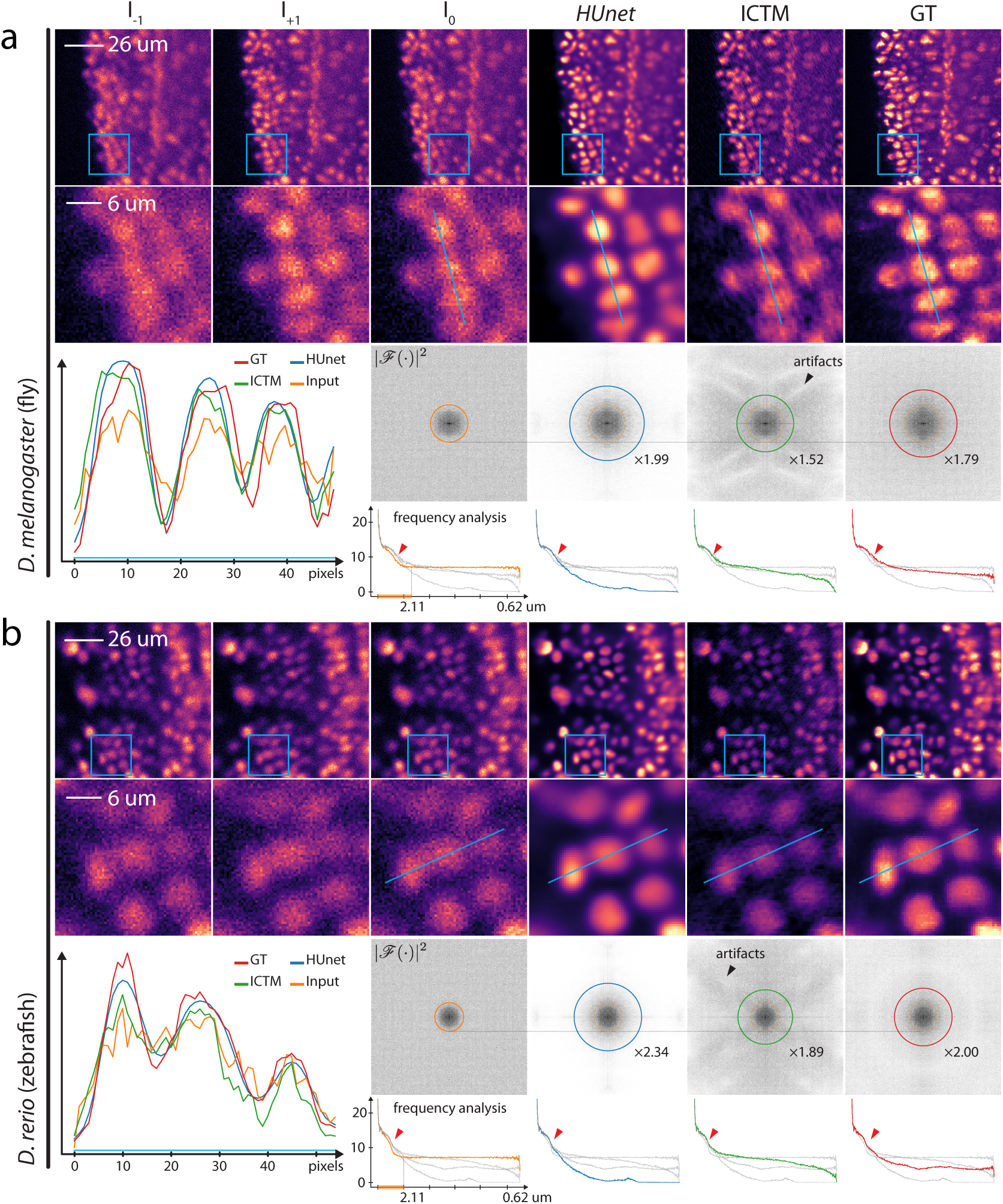
Comparison between phase-diversity based *HUnet* deconvolution (3 image input *I*−_1_, *I*_0_, *I*_+1_), non-blind *ICTM* deconvolution (*I*_0_ as input), and ground truth, for both (a) *D. melanogaster* (fly) and (b) *D. rerio* (zebrafish) embryo images. In addition, line profiles and Fourier frequency analysis are shown for aiding comparison of results. Quantitatively, *HUnet* deconvolved images have a wider frequency (2.3 fold) support and a low noise floor compared to the input images (according to Mizutani’s single image resolution estimate [29]).

### Spatially-variant blind deconvolution

In-toto live imaging of developing embryos involves imaging large fields of view of up to 800 µm. Almost certainly, the sample’s varying refractive index composition is not constant over the field of view [30] which leads us to consider the problem of deconvolving images with spatially varying optical aberrations. Tackling spatial variance is challenging for standard wavefront correction approaches because corrections are necessarily limited to a small region of the field of view [30]. In contrast, blind deconvolution – possibly informed by phase information – can address this problem. First, wee simulate spatial variance by varying the PSF over tiles of dimensions 32 × 32. At inference time, for *Znet*, we simply break large images into smaller tiles (256 × 256). For *HUnet* we rely on translation invariance. The results are shown in Fig. 4. Deconvolution with *HUnet* produces sharper outputs with overall 33% wider frequency support. Similarly, *Znet* predictions seem to match the expected behaviour of predicting the average PSF over the tile – however, in some cases (yellow inset) the predicted PSF is incorrect. These results suggest that for estimating spatially varying aberrations, training should be best done directly on a variant dataset.

**Fig. 4.**
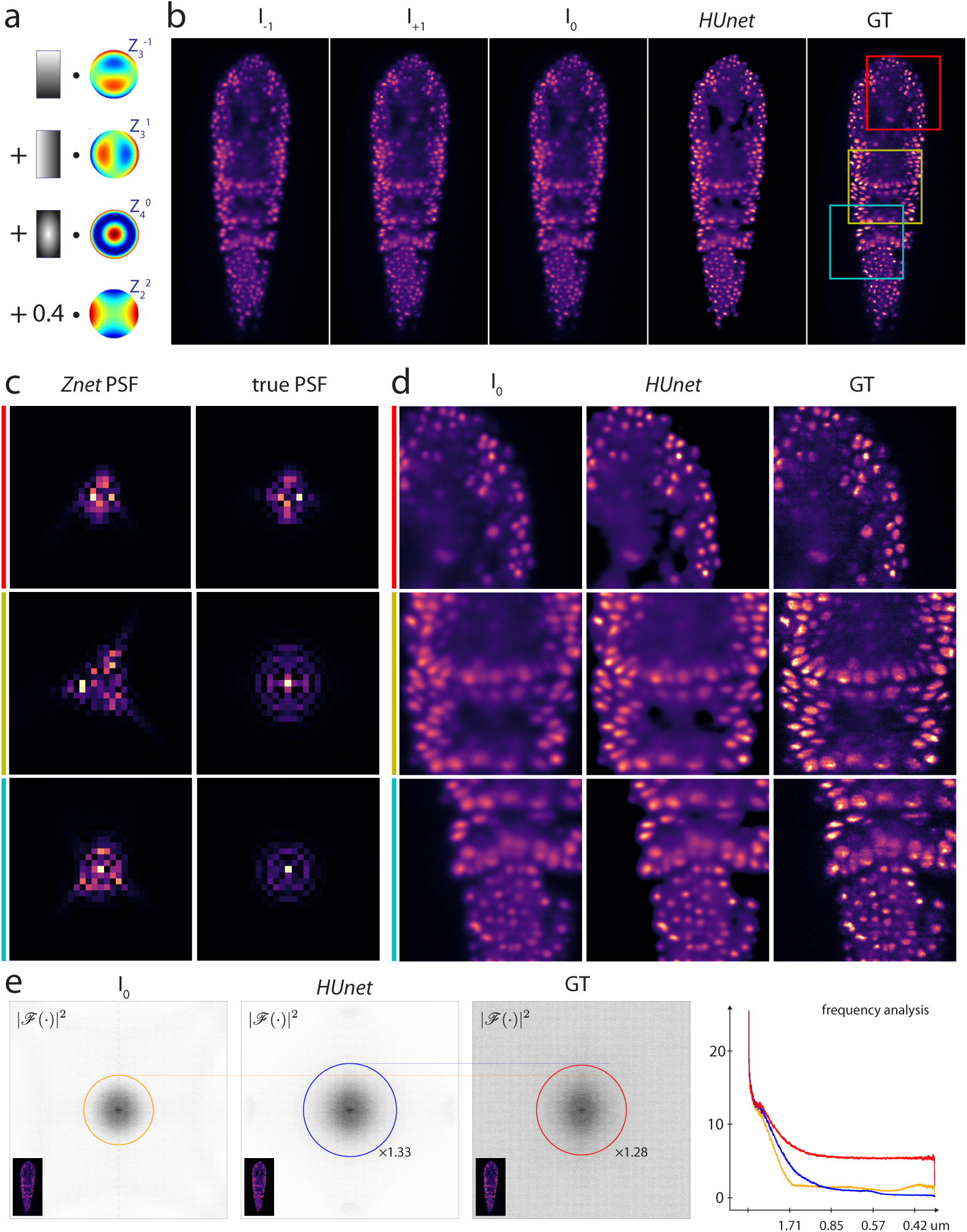
Performance of *HUnet* and *Znet* for spatially varying aberrations. **(a)** Pictorial description of the spatially variant aberrations used for evaluation. Here we have the sum of primary spherical 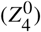 varying with a quadratic profile, vertical 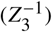 and horizontal 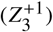 coma varying linearly along Y and X, respectively, and constant astigmatism 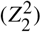. **(b)** phase-diversity stack (*I*_− 1_ *I*_+1_, and *I*_0_) used as input. **(c)** Predicted versus true PSFs. The PSFs shown here are the average of the spatially varying PSFs within each tile. **(d)** Expanded images for insets in **(a)** with the red inset along row 1, yellow along row 2 and cyan along row 3. **(e)** Frequency analysis shows that *HUnet* deconvolution frequency support is comparable to that of the ground truth, only with less noise at high frequencies.

### Importance of phase diversity

We found very similar deconvolution performance for *HUnet* trained with and without phase diversity (SSIM reduction of 0.017 for fly and 0.010 for zebrafish, see Table 1) suggesting that the implicit phase information is either not used or not necessary to achieve deconvolution. Our interpretation is that the implicitly learned image prior – cell nuclei – is strong enough, rendering the phase information mostly superfluous. This encouraging result suggests that for moderate optical aberrations, blind deconvolution of stereotypical images does not require phase information. This is in agreement with previous results on isotropy restoration [18, 31] that showed that axial deconvolution is possible in the absence of phase information for images that have strong stereotypy. However, for more severe aberrations and noisier acquisitions, phase retrieval – as proposed here or otherwise – in concert with wavefront correction is likely required for good results.

### Aberration estimation

The phase estimation model *Znet* performed reliably in predicting astigmatism 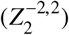, coma and trefoil aberrations 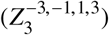, as well as spherical aberration 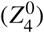, but less so for quadrafoil and secondary astigmatism (Fig. 5). In contrast to the deconvolution model (*HUnet*), the aberration estimation model (*Znet*) trained on fly images generalized better to zebrafish images (5% vs 11% performance reduction) suggesting less reliance on nuclei image priors and actual utilisation of the phase diversity information (Table 3). As expected, and in contrast to our deconvolution model, aberration estimation (*Znet*) models trained without phase-diversity completely fail to generalise to the test set. In the best case, these models over-fit to the training set in the absence of dropout regularisation.

**Table 3.** *Znet* performance. Training dataset along rows and testing dataset along columns. Two *Znet* architectures with 14 and 19 million parameters where tested.

|  |  | fly (test) |  | zebrafish (test) |  |
| --- | --- | --- | --- | --- | --- |
| | | $r^2$ | MAE | $r^2$ | MAE |
| 14M | fly | 0.739 | 0.134 | 0.791 | 0.124 |
|  | zebrafish | 0.622 | 0.163 | 0.841 | 0.106 |
| 19M | fly | 0.738 | 0.136 | 0.786 | 0.125 |
|  | zebrafish | 0.576 | 0.176 | 0.838 | 0.107 |

**Fig. 5.**
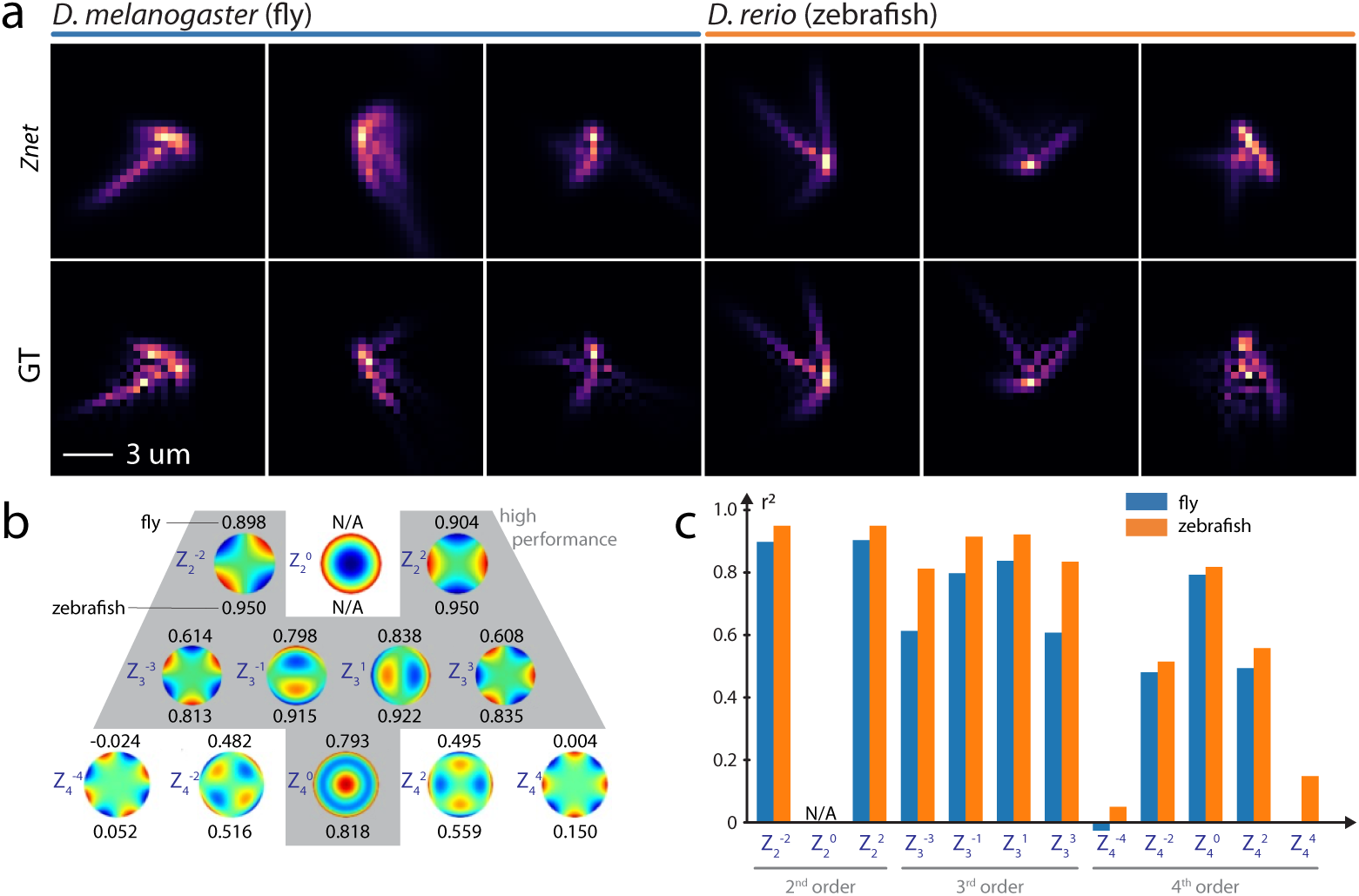
**(a)** *Znet* Example PSF predictions versus corresponding ground truth PSFs. **(b)** *Znet* model performance per coefficient on fly (top) and zebrafish (bottom) datasets. The coefficient of determination (*r*^2^, within [0, 1]) is given for each Zernike coefficient. Gray background area identifies coefficients for which model performance is high (above 80%). Defocus is not considered as we assume in-focus imaging. **(c)** Bar chart showing that Zernike coefficients with high angular meridional frequency are difficult to predict – for example 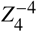 and 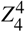.

### Application to light-sheet acquisitions

After promising results on simulations, we wanted to test our deconvolution model (*HUnet*) on images acquired on a light-sheet microscope – with all the imperfections and non-idealities that this entails. In light of these circumstances, our results are promising: Fig. 6 shows an overall 46% increase in resolution as well as suppression of all high-frequency noise – a typical and beneficial side-effect of CNN inference. Yet, for deeper imaging regions (lower panels in Fig. 6) our model only marginally improves image quality because of the limits of our assumptions. Indeed, our simulation model implicitly assumes (i) an illumination confined to an infinitely thin light-sheet and (ii) no scattering on both illumination and detection paths. While overall performance is satisfactory for images taken at shallow imaging depths within the zebrafish embryo, for deeper regions, we observe strong out-of-focus light and background due to a thick light-sheet and detection scattering – the resulting images, while apparently sharper, should be treated with caution because the input images strongly violate our assumptions. Overall, our observations reinforce the notion that low-order aberrations (as corrected by *HUnet*) are not the only – or even dominant – cause of image quality degradation in light-sheet imaging of large samples such as zebrafish embryos [30]. Instead, higher-order aberrations and scattering on both illumination and detection paths account for most of the image quality degradation.

**Fig. 6.**
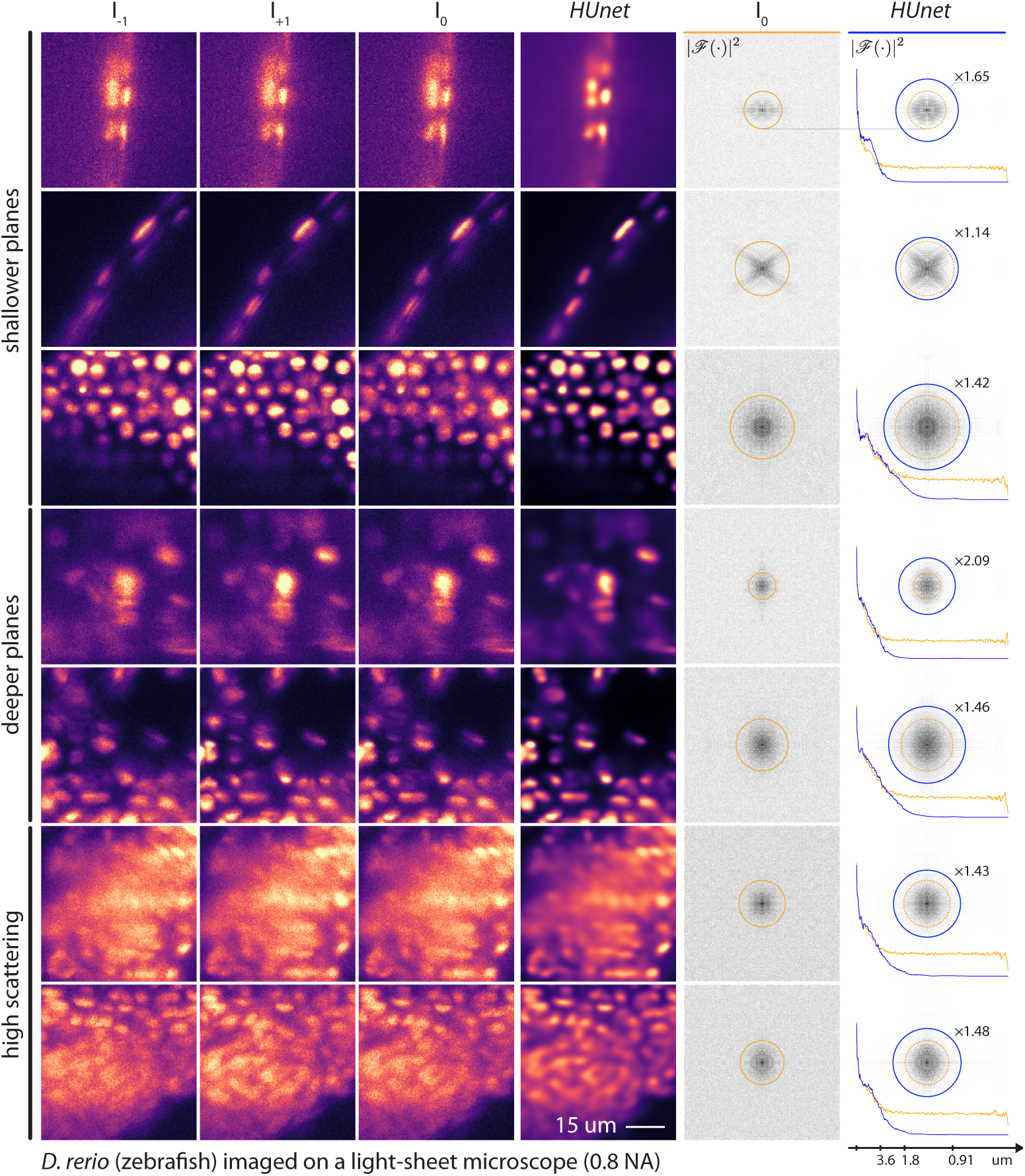
Applying HUnet deconvolution on phase-diverse acquisitions from a light-sheet microscope (0.8 NA, 16×, pixels 0.406). Applying *HUnet* to input images *I*_−1_, *I*_0_, and *I*_+1_ results in sharper images as evidenced by the Fourier spectra comparison between input *in-focus* image *I*_0_ and the *HUnet* image. Frequency analysis suggests that *HUnet* images have a 46% wider frequency support (median value for all 7 images). Moreover, *HUnet* expectedly suppresses high-frequency noise and in-paints missing frequencies to achieve sharper nuclei boundaries.

### Varying network size for *Znet*

We investigated whether increasing the phase estimation network size from 14 to 19 million parameters could further improve results. Our results show the same overall performance. The larger network performance on the fourth order coefficients was slightly reduced in the fly dataset and marginally better on the zebrafish dataset with similar performance on the second and third order coefficients (Fig. 5). However the generalization of the larger network trained on zebrafish embryo images was reduced when applied to fly embryo images – suggesting over-fitting to dataset specific features (Table 3).

### To train jointly, or not

Unfortunately, we could not find a configuration in which joint training would significantly improve the performance of *HUnet* for both datasets. While it improves *Znet* performance slightly for zebrafish dataset, it reduces performance for fly dataset. Possible explanations are: (i) using a trained *PSFnet* instead of a differentiable analytical model is sub-optimal since a network can only approach and not attain the exactness of an analytical model, (ii) the consistency loss only refers to *I*_0_ instead of to the whole phase diversity stack *I*_−1_, *I*_0_, and *I*_+1_, which likely would provide stronger consistency information, (iii) the large mismatch between the number of parameters in *HUnet* and *Znet* (Table 4), (iv) finally, the non-ideal nature of self-consistency loss: the mean absolute error on blurred images might not be helpful. Overall, more work is needed to understand how and in which circumstances physics based consistency losses could help.

**Table 4.** Joint training performance. Training dataset along rows and testing dataset along columns.

|  |  | fly (test) |  | zebrafish (test) |  |
| --- | --- | --- | --- | --- | --- |
| | | $r^2$ | SSIM | $r^2$ | SSIM |
| in-focus |  | 0.691 | 0.264 | 0.869 | 0.475 |
| *HUnet* | fly | 0.925 | 0.416 | 0.971 | 0.750 |
|  | zebra | 0.920 | 0.464 | 0.974 | 0.891 |
| | | $r^2$ | MAE | $r^2$ | MAE |
| *Znet* | fly | 0.688 | 0.148 | 0.742 | 0.140 |
|  | zebra | 0.581 | 0.168 | 0.844 | 0.103 |

## 5. Discussion

We have shown on simulations that convolutional models can be trained to deconvolve (*HUnet*) and estimate aberrations (*Znet*) from simulated phase diverse acquisitions. Moreover, our experiments with images from a light-sheet microscope also confirm improved resolution (46% wider frequency support) even in regions where assumptions made during training are not fully guaranteed – i.e. in the presence of confounding factors such as high-order aberrations, scattering and variance in the light-sheet thickness.

The most interesting observation is the robust performance of our deconvolution model (*HUnet*) in the absence of phase-diversity. This suggests that the model can directly be used for blind deconvolution of single images. How is this possible? Our interpretation is that the model captures an implicit prior of the highly stereotypical structure of fluorescently labelled nuclei and does not require phase information to be competitive. This suggests that such models can be made quite robust against uncertain aberration regimes as long as the aberrations are not too severe. Of course, such trained models are necessarily limited to the stereotypy of their training data.

In contrast, and as expected, phase estimation via *Znet* does require phase-diverse images as input. With the caveat that a direct comparison of models trained on different datasets is non-ideal, our phase estimation model outperforms the semi-blind model in [12]. *Znet* achieved a mean *r*^2^ of 0.74 and 0.84 on fly and zebrafish datasets respectively for regressing astigmatism, coma and trefoil – numbers to be compared to a *r*^2^ of 0.58 in [12] for regressing focus and astigmatism. The reduced performance on the fourth order coefficients could be attributed to potential aliasing of the PSF.

Future work could focus on technical aspects such as investigating how to improve performance with adversarial loss training, optimising the architecture for single image inputs, as well as improving our simulation model. Moreover, it is possible to replace *PSFnet* with a non-learned differentiable analytical function. More importantly, extending our framework in 3D would be key to be able to deconvolve acquisitions axially.

From an adaptive optics perspective, the strong performance of *Znet* suggests that it could be used as part of an online adaptive optics setup for the purpose of closed-loop wavefront estimation and correction. In this context, the advantage of using neural network inference is its speed compared to iterative schemes – it runs a single pass per input and is often hardware accelerated. However, it is unclear that defocus-only phase-diversity *per se* is the right approach due to the lack of contrast between the images within the stack. Another approach, perhaps involving complex phase relations like those in spiral phase masks, may provide better contrast [32], and would thus provide more robustness to noise and better performance for higher order aberrations.

From a light-sheet imaging perspective, our experimental evidence suggests that low-order optical aberrations are not the sole, and perhaps not the dominant, cause of image degradation. As we have shown, scattering on both illumination and detection paths offer considerable obstacles to methods that assume a thin light-sheet, low-order aberrations, and only ballistic photons. It would interesting to see what deep learning can do to address these issues.

## Acknowledgments

We thank Dr. Jan Huisken from the *Morgridge Institute for Research*, Madison, Wisconsin, USA for kindly providing zebrafish transgenic lines (*h2afva:h2afva-mCherry*). Thanks to our *Chan Zuckerberg Biohub* colleagues Li-Hao Yeh and Shalin Metha for discussion, advice and feedback. Finally, we thank the *Chan Zuckerberg Biohub* and its donors for funding this work.

